# A high-quality genome assembly and annotation of the European earwig *Forficula auricularia*

**DOI:** 10.1101/2022.01.31.478561

**Authors:** Upendra R. Bhattarai, Mandira Katuwal, Robert Poulin, Neil J. Gemmell, Eddy Dowle

## Abstract

The European earwig *Forficula auricularia* is an important model for studies of maternal care, sexual selection, sociality and host-parasite interactions. However, detailed genetic investigations of this species are hindered by a lack of genomic resources. Here we present a high-quality hybrid genome assembly for *F. auricularia*. The genome was assembled using nanopore long-reads and 10x chromium link-reads. The final assembly is 1.06Gb in length with 31.03% GC content. It consists of 919 scaffolds with an N50 of 12.55Mb. Half of the genome is present in only 20 scaffolds. Benchmarking Universal Single-Copy Orthologs scores are ~90% from three sets of single-copy orthologs (eukaryotic, insect, and arthropod). The total repeat elements in the genome are 64.62%. The MAKER2 pipeline annotated 12,876 protein-coding genes and 21,031 mRNAs. A phylogenetic analysis revealed the isolate used in our genomic analysis belongs to Subspecies B, one of the two known genetic subspecies of *F. auricularia*. The genome assembly, annotation, and associated resources will be of high value to a large and diverse group of researchers working on Dermapterans.

## Introduction

Insects have been at the forefront of genetic research for various biological questions (Wilson-Sanders 2011; Mukherjee *et al.* 2015; Simons and Tibbetts 2019). However, most of the genetic studies are carried out on a small number of holometabolous insects that undergo true metamorphosis. In contrast to Holometabola, hemimetabolous insects undergo incomplete metamorphosis with a series of nymphal moults that increasingly resemble the adult form (Truman 2019). It is widely accepted that Holometabola branched out from hemimetabolous ancestors during the Permian 300 Mya (Labandeira and Phillips 1996; Yang 2001). Yet the conserved mode of development, embryonic organisation, and the adult body plan of hemimetabolous insects offer a unique model for the study of developmental and evolutionary mechanisms. However, even with the increasing number of sequenced genomes, the majority belong to the Holometabola (Ylla *et al.* 2021). This has been a bottleneck for the exploration of the diverse biology and life history of hemimetabolous insects. To address this paucity, we report a high-quality annotated genome of the European earwig, *Forficula auricularia* (Dermaptera: Forficulidae).

The European earwig *Forficula auricularia* is widely distributed, comprising two recognised subspecies, A and B (Wirth *et al.* 1998). They are native to the western Eurasian region and were introduced to North America, Australia, and New Zealand, where they have quickly adapted and became abundant throughout the regions (Quarrell *et al.* 2018; Tourneur and Meunier 2019). Their propensity to dwell on flower and kitchen gardens can cause significant damage to crops, flowers, and commercial vegetables and make them important agricultural pests (Campos *et al.* 2011; Hill *et al.* 2019).

They have been of particular interest for many researchers not just because of their importance in the agricultural ecosystem (Binns *et al.* 2021) but also their importance as a research model for various biological and evolutionary phenomena like sexual selection, maternal care, family interactions, reproductive strategy, and social behaviour (Forslund 2000; Falk *et al.* 2014; Kramer *et al.* 2015; Van Meyel and Meunier 2020). They have been extensively studied by behavioural ecologists for the early evolution of group-living and family life (Falk *et al.* 2014). The male earwigs also show an unusual bias in their use of lateral left and right sexual organs without any conspicuous anatomical differentiation (Kamimura 2006). Like the right-handedness in humans, 90% of males of giant earwig *Labidura riparia* show a preference for the right penis for copulation, providing insights into the evolutionary origin of lateralization (Kamimura *et al.* 2021). Similarly, they are an excellent lab model to study extended phenotypes as they exhibit strange suicidal water-seeking behaviour during the late stages of infection by mermithid nematodes (Herbison *et al.* 2019). However, their use as a genetic model has been severely limited by the lack of a reference genome.

Here, we have sequenced, assembled, annotated, and analysed the genome of the European Earwig, *Forficula auricularia*, and confirmed the subspecies identity of the individuals we used. This genome will help researchers study multiple facets of this insect’s exciting biology and evolutionary characters and broaden our understanding of insect and genome evolution.

## Methods and Materials

### Sample collection and preparation

Earwigs (*Forficula auricularia*) were collected from the Dunedin Botanic Garden (−45° 51’ 27.59” S, 170° 31’ 15.56” E) and reared in a temperature-controlled room (Temperature: cycling from 15 to 12°C, day/night; Photoperiod of L:D 16:8) in the Department of Zoology, University of Otago, Dunedin. Earwigs were snap-frozen in liquid nitrogen and stored at −80°C before dissection and subsequent nucleotide extraction. Earwigs were dissected in 1x PBS buffer under a dissection microscope to check for nematode parasites, and only non-parasitized individuals were used in this study. The head, wings and muscles from the thorax region were used for DNA extraction. Juvenile instars required for RNA extraction were obtained directly from the field.

### DNA extractions

DNA was extracted using either the Nanobind Tissue Big DNA kit (Circulomics, USA) or DNeasy Blood & Tissue Kit (Qiagen, Germany) by following the manufacturer’s protocol. Tissues from a single individual were used for each extraction. After the extraction, RNase treatment was performed using 4μl of RNase A (10mg/ml) per 200μl of DNA elute. DNA was quantified in a Qubit 2.0 Fluorometer (Life Technologies, USA) and quality analysed using Nanodrop. Low quality DNA samples were further cleaned with 1.8x by volume AMPure XP beads (Beckman Coulter, USA), wherever applicable, following the manufacturer’s instructions and eluted in 55μl of molecular grade water. High-quality DNA samples were stored at −20°C and were used within a week of extraction.

### Linked read library preparation and sequencing

Linked read library was prepared at the Genetic Analysis Services (GAS), University of Otago (Dunedin, New Zealand). DNA from an adult male was extracted using the Nanobind kit and size-selected for fragments over 40kbp using Blue Pippin (Sage Science, USA). A chromium 10X linked read (10× Genomics, USA) library was prepared following the manufacturer’s instructions. The library was sequenced on the Illumina Nova-seq platform to generate 2×151bp paired-end reads (Garvan Institute, Australia).

### Long read library preparation and sequencing

Five long-read sequencing libraries for Oxford minion were prepared using a ligation sequencing kit (SQK-LSK109) (Oxford Nanopore Technologies, Oxford, UK) following the manufacturer’s instructions. To increase the raw nanopore read N50 the first and the second libraries were prepared using 1.75 and 0.75μg of DNA extracted via a Circulomics kit from two adult male earwigs. Both libraries were sequenced in a single minion flow cell, flushing the flow cell to remove remains of the first library before loading the second library with a Flow cell wash kit (Oxford Nanopore Technologies, Oxford, UK).

To increase the total raw output the third and the fourth libraries were prepared with DNA from two adult female earwigs, both extracted with a Qiagen kit followed by the AMPure XP beads clean-up step. Input DNA for these two libraries were 2.6 and 3.2 μg. These were each sequenced on an individual minion flow cell. The fifth library was prepared using 3.0 μg of DNA from an adult male earwig. As before DNA was extracted using a Qiagen kit followed by AMPure XP beads clean-up. However, before library preparation, the DNA was sheared five times using a 26G X 0.5” needle (Terumo, Japan). All prepared libraries were sequenced with R9 chemistry MinION flow cell (FLO-MIN106) (Oxford Nanopore Technologies, UK).

### RNA extraction and sequencing

Total RNA from the different developmental stages, sex, and tissues was extracted using a Direct-zol RNA MicroPrep kit (Zymo Research) with an on filter DNAse treatment following the manufacturer’s instructions. Samples included: whole body (gut removed) of juvenile instars 1-2 and juvenile instars 3-4, dissected tissues (antennae, head, thorax, abdomen, legs, and gonads) of adult males and females. RNA from each individual and tissue type was extracted separately. RNA was quantified on a Qubit 2.0 Fluorometer (Life Technologies, USA) and initially quality checked using a nanodrop. Only high-quality extracts were further processed and were stored at −80°C until use.

RNA integrity was evaluated on a Fragment Analyzer (Advanced Analytical Technologies Inc., USA) at the Otago Genomics Facility (OGF), University of Otago, Dunedin, New Zealand. As with most of the insect RNA extracts (Winnebeck *et al.* 2010) RNA quality number (RQN) values ranged from 2.5 to 10 due to the collapsing of the 28S peak; quality was thus determined via the trace rather than RQN. Four pools of samples at equimolar concentration underwent library preparation. Pools consisted of: 8 whole body extractions for juvenile instar 1-2, 8 whole body extractions for juvenile instar 3-4, individual body tissues from five adult males, and individual body tissues from five adult females. TruSeq stranded mRNA libraries were prepared and sequenced as 2×100bp paired-end reads across two lanes of HiSeq 2500 Rapid V2 flowcell at the OGF.

### Genome size estimation

Flow cytometry and k-mer based approach with short-read data were used to estimate the genome size. Flow cytometry analysis was performed on a single head of earwig with two biological replicates at Flowjoanna (Palmerston North, NZ). Briefly, the earwig’s head was dissociated with a pestle in 500µl of the stock solution containing 0.1% w/v trisodium citrate dihydrate, 0.1% v/v IGEPAL, 0.052% w/v spermine tetrahydrochloride, and 0.006% sigma 7-9 (all Sigma-Aldrich, USA). Rooster red blood cells (RRBC) derived from the domestic chicken (*Gallus gallus*), stored in citrate buffer, was used as a reference sample. Test samples were filtered through 35µl filter cap and further dissociated by adding 100µl of 0.21mg/ml trypsin followed by 75µl of 2.5mg/ml trypsin inhibitor (both Sigma-Aldrich) for 10 minutes at 37°C. Nuclei were stained using 100µl of pre-stain (containing 416mg/ml propidium iodide (PI) with 500mg/ml RNAse in-stock solution). Two sample tubes, one prepared with RRBC and one prepared without, were then processed on a FACSCalibur (BD Biosciences, USA). The instrument was equipped with a 488nm laser to produce fluorescence collected using the FL-2-Area signal (585/42 BP), along with forward scatter (FSC) and side scatter (SSC) signals that enabled RRBC nuclei to be resolved from earwig nuclei. Data were analysed using Flowjo (BD Biosciences, USA) and the pg/nuclei of the sample calculated.

For k-mer based genome size estimation, we used the paired-end linked read sequences. Reads were processed with the scaff_reads script from Scaff10x (v.5.0) to remove the 10x link adapters. Quality control was carried out with Trimmomatic (v.0.39) (options: SLIDINGWINDOW:4:15 LEADING:5 TRAILING:5 MINLEN:35). We used KMC (v.3.1.1) (Kokot *et al.* 2017) with a k-mer size of 21 to produce a histogram, which was then visualised in Genomescope (v.2.0) web browser.

### Bioinformatic pipeline

All the scripts used for genome assembly, *denovo* repeat library construction, and annotation are available on GitHub (https://github.com/upendrabhattarai/Earwig_Genome_Project). The bioinformatics software and packages were run in New Zealand eScience Infrastructure (NeSI). Below is a description of the pipeline (Figure 1).

**Figure 1:**
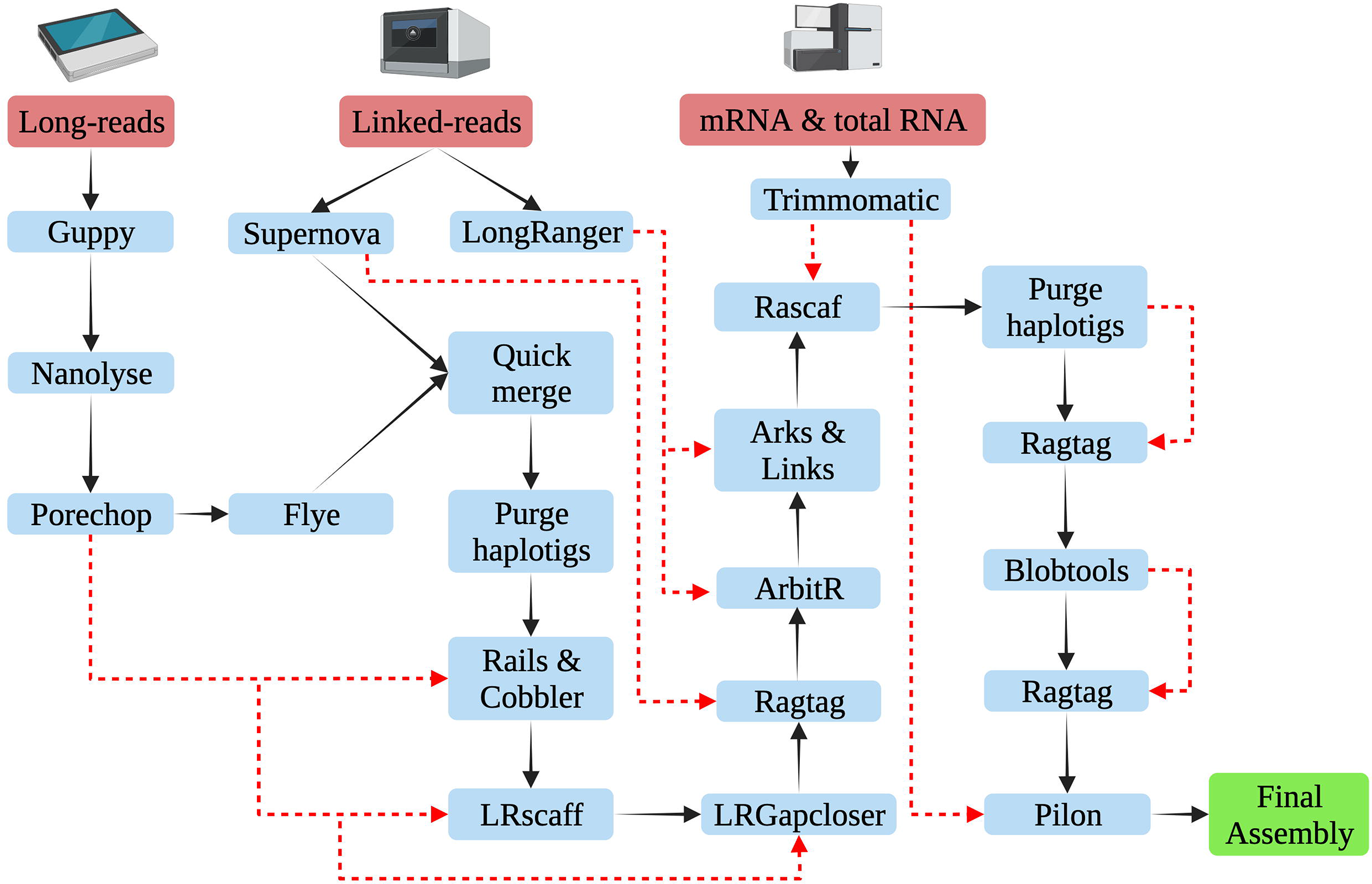
Schematic representation of the assembly pipeline for the *Forficula auricularia* genome. The black arrow represents the workflow and the red dotted lines represent the additional input data in the pipeline (Created with BioRender.com).

### Genome assembly

Paired-end Illumina reads from the chromium library were assembled using Supernova (v.2.1.1) (Weisenfeld *et al.* 2017). Assembly quality was assessed using Quast (Gurevich *et al.* 2013) and BUSCO (Simão *et al.* 2015) metrics such as N50 values, contig/scaffold number, contiguity and ortholog completeness. Based on several trial assemblies, we down-sampled the total input to 660 million paired-end reads to produce an assembly with better completeness and contiguity. The assembled fasta sequence was obtained with “pseudohap” style of the supernova “mkoutput” function.

Nanopore reads were basecalled using guppy (v.5.0.7) and assembled using Flye (v.2.7.1) (Kolmogorov *et al.* 2019) with default parameters. The assembly statistics (as mentioned above) for the Flye assembly were better than that for the Supernova assembly. Therefore, the supernova assembly was merged into the Flye assembly using Quickmerge (Chakraborty *et al.* 2016). The resulting assembly was processed with Purgehaplotigs (Roach *et al.* 2018) to remove redundant and duplicated contigs.

The purged genome underwent further scaffolding and gap-closing steps using Rails (v.1.5.1) and Cobbler (v.0.6.1) (Warren 2016), Lrscaf (v.1.1.11) (Qin *et al.* 2019) and Lrgapcloser (Xu *et al.* 2019) with the raw long-read data. The resulting assembly was scaffolded with Ragtag (v.2.1.0) (Alonge *et al.* 2021) using the Supernova assembly. The raw linked-read data were further used to scaffold the assembly with ArbitR (v.0.2) (Hiltunen *et al.* 2021), Arks (v.1.0.4) (Coombe *et al.* 2018) and Links (v.1.8.7) (Warren *et al.* 2015). mRNA-seq reads sequenced for genome annotation purposes, and total RNA-seq reads sequenced for another project (manuscript under preparation) were also used for scaffolding the assembly with the Rascaf (Song *et al.* 2016). Duplicated and redundant haplotigs were again removed using Purgehaplotigs, and discarded haplotigs were used for scaffolding the assembly using Ragtag. Blobtools2 (Laetsch and Blaxter 2017) was used to remove small (<1000bp) and low coverage contigs (<5x coverage). Contigs that were filtered out were used for re-scaffolding the assembly with Ragtag. Finally, we used Pilon (v.1.24) to polish the assembly using mRNA-seq data.

### Repeat content analysis

To assist with annotation a custom repeat library was generated for the Earwig genome using different *de novo* repeat and homology-based identifiers, including LTRharvest (Ellinghaus *et al.* 2008), LTRdigest (Steinbiss *et al.* 2009), RepeatModeler (Flynn *et al.* 2020), TransposonPSI (Brian Haas 2010) and SINEBase (Vassetzky and Kramerov 2013). We concatenated the individual libraries, and sequences with more than 80% similarity were merged to remove redundancy using usearch (v.11.0.667) (Edgar 2010). It was then classified using RepeatClassifier. Sequences with unknown categories in the library were mapped against the UniProtKB/Swiss-Prot database (e-value <1e-01); if sequences were not annotated as repeat sequences they were removed from the library. The final repeat library was used in RepeatMasker (v.4.1.2) (Chen 2004) to generate a report for genome repeat content and provided to the Maker2 pipeline to mask the genome.

### Genome annotation

Genome annotation was carried out with three iterations of the MAKER2 (v.2.31.9) (Holt and Yandell 2011) pipeline combining evidence-based and ab initio gene models. The first round of Maker used evidence-based models and the other two rounds were run using ab initio gene models. For the first round, we provided the MAKER2 pipeline with 180,119 mRNA transcripts *denovo* assembled via the Trinity pipeline (Grabherr *et al.* 2011) along with 26,414 mRNA and 1,529 protein sequences of dermapterans from NCBI and 779 dermapteran protein sequences from the Uniprot database.

Augustus was trained using Braker (v.2.16) (Hoff *et al.* 2019) and SNAP was trained after each round of MAKER to use for ab initio gene model prediction. For the functional annotation, we ran InterProScan (v.5.51-85.0) (Jones *et al.* 2014) for the predicted protein sequences obtained from MAKER and retrieved InterPro ID, PFAM domains, and Gene Ontology (GO) terms. Furthermore, we ran BLAST with the Uniprot database to assign gene descriptors to each transcript based on the best BLAST hit.

### Phylogenetic analysis

Two sibling species of *F. Auricularia* have been described (Wirth *et al.* 1998). To assess which of these we sequenced, nucleotide sequences covering the COI and COII region from 34 isolates of *F. auricularia* were downloaded from NCBI. Those included 15 isolates reported by Wirth et al. (1998) originally used to infer sibling species A and B and other isolates from Belgian orchards submitted to NCBI. Nucleotide sequence covering COI and COII regions from the assembled genome was extracted through BLAST hits. The same genomic region extracted from the mitochondrial genome of *Euborellia arcanum* was used as an outgroup. Nucleotide sequences were aligned using Clustal Omega (v1.2.3). The evolutionary history was inferred using the Neighbor-Joining method (Saitou and Nei 1987) in the bootstrap test of 1000 replicates (Felsenstein 1985). The evolutionary distances were computed using the Maximum Composite Likelihood method (Tamura *et al.* 2004) and are in the units of the number of base substitutions per site. All ambiguous positions were removed for each nucleotide sequence pair (pairwise deletion option). There were a total of 799 positions in the final dataset. The optimal tree is shown (Figure 5) and the evolutionary analyses were conducted in MEGA11 (Tamura *et al.* 2021).

## Results and Discussion

### Genome size estimates

The flow cytometer estimated the genome size of 968.22±20.747 Mb (Mean±SD) for the earwig genome. Similarly, the kmer based approach using adapter removed paired-end data from linked read sequencing estimated the male earwig to be 988 Mb. Whereas, an earlier estimation of an unknown dermapteran (earwig) species genome size was 1.4 Gb (Gregory 2005) showing a variable genome size within the order.

### Genome assembly

A total of 799.6 million paired-end reads was generated for the linked-read library. Downsampled to 660 million paired-end reads, Supernova estimated the genome size of 1.22 Gb, raw coverage of 82.02%, effective coverage of 39.50% and weighted mean molecule size of 22.45 kb. The Supernova assembly was 1.15 Gb in size and had 145,055 contigs, with an N50 of 0.03 Mb and L50 of 7,500. Quast reported a complete BUSCO of 64.69% and a partial BUSCO of 9.24% from the eukaryotic database.

The nanopore sequencing yielded approximately 10.7 Gb of data, consisting of over 3 million reads. The median read length was 897 bp with an N50 length of 11,986 bp (Supplementary Table 1). The median read Phred quality was 13.34. Primary assembly from Flye produced an assembly of 1.1 Gb. There were 187,66 contigs with N50 of 0.18 Mb and L50 of 1,832. Quast reported a complete BUSCO of 82.18% and a partial BUSCO of 9.24%. The long-read assembly showed better contiguity and completeness, so we merged the linked-read assembly with the long-read assembly (Table 1).

**Table 1.** Assembly statistics for the genome assembly of the European earwig *Forficula auricularia.* Quast scores are to its default Eukaryota database.

|  | Assembly length | No. of scaffolds | N50 | L50 | Ns per 100 kbp | BUSCO % (Quast) |  |
| --- | --- | --- | --- | --- | --- | --- | --- |
|  |  |  |  |  |  | Complete | Partial |
| Supernova assembly | 1,145,470,221 | 145,055 | 30,358 | 7,500 | 3,677.89 | 64.69 | 9.24 |
| Flye assembly | 1,118,374,848 | 18,766 | 180,737 | 1,832 | 0.35 | 82.18 | 9.24 |
| Final hybrid assembly | 1,062,210,345 | 919 | 12,548,649 | 20 | 846.85 | 87.13 | 2.97 |

The final hybrid assembly has a size of 1.06Gb. It has 919 scaffolds with an N50 of 12.55Mb, which shows that the assembly is highly contiguous. Half of the genome is present in just 20 scaffolds, as denoted by the L50 number (Table 1). Assembly has 846.85 “N’s” per 100kbp. The BUSCO score from the insect database (n=1,367) for the assembly was 87.1% complete, among which 4.1% were duplicated, and 3.1% fragmented BUSCO (Figure 2). Improvement in assembly statistics after each processing step is given in Supplementary Table 2.

**Figure 2.**
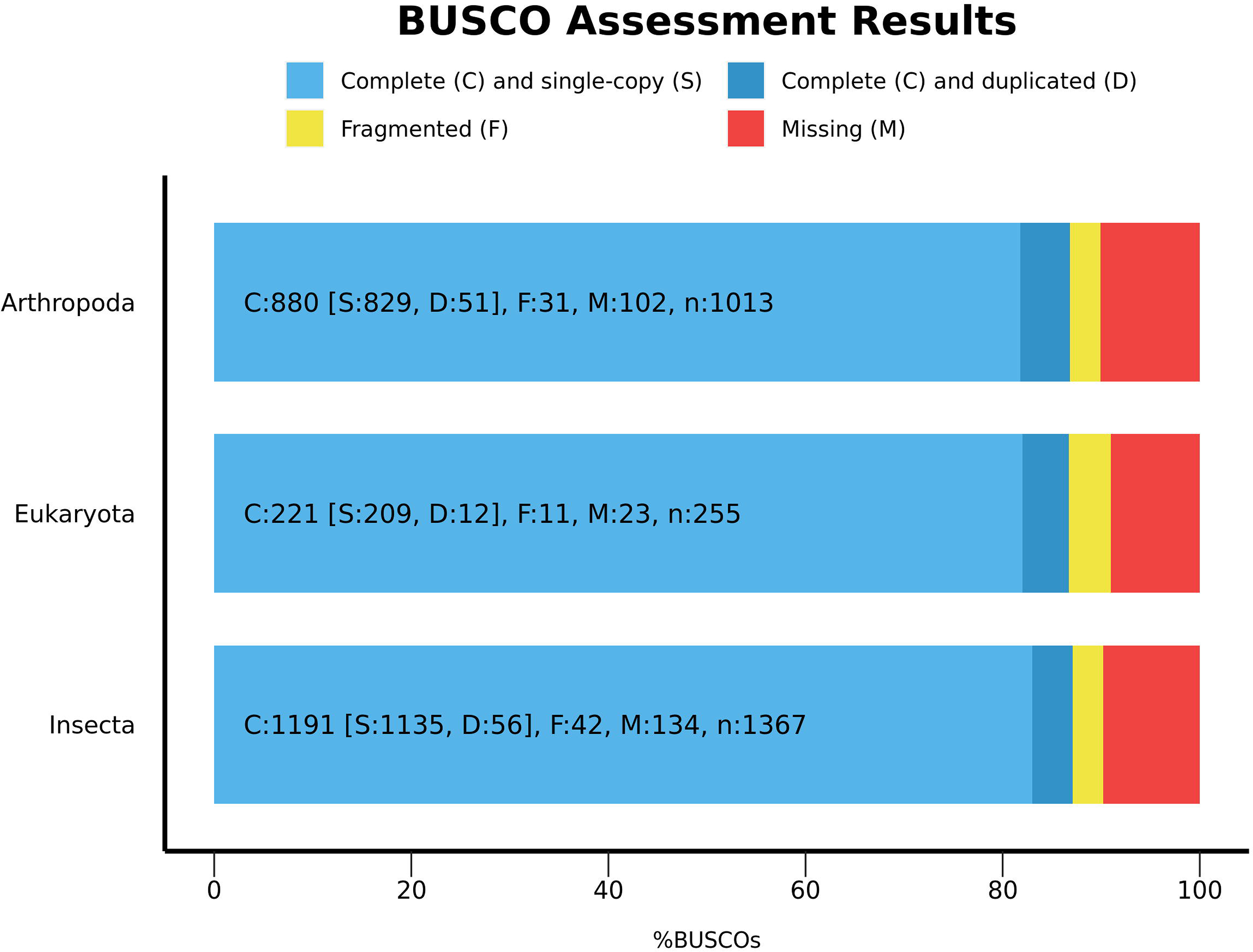
The BUSCO (v.5) report for the final hybrid assembly of the *Forficula auricularia* genome. BUSCO scores in percentage (x-axis) from Arthropoda, Eukaryota, and Insecta (odt10) databases (y-axis) are shown in the bar plot. The light blue portion of the bar represents Complete and single-copy orthologs, dark blue represents complete and duplicated orthologs, yellow represents fragmented BUSCO genes and red represents missing BUSCO genes.

The only other whole-genome sequence publicly available from the Dermaptera order is of the other earwig *Anisolabis maritima*. Using the insect database (n=1,367) it has 83.4% complete and 10.8% fragmented BUSCO scores. The *A. maritima* assembly has an N50 of 1.4Mb with a genome size of 649.7 Mb (Mei *et al.* 2022). So, in comparison, the *F. auricularia* genome assembly has a better gene model and contiguity from the Dermaptera order.

### Genome repeat contents

Repeat analysis of the assembly showed that interspersed repeats comprised ~686 Mb (64.62%) of the *F. auricularia* genome. This includes ~248 Mb of retroelements (23.37% of the genome), ~178 Mb of DNA transposons (16.79% of the genome), ~35 Mb of rolling-circles (3.28% of the genome) and ~260 Mb of unclassified elements (Table 2). Unusually large and variable genome sizes characterise Hemimetabolans (Wu *et al.* 2017). Comparative analysis in six species of Gomphocerine grasshoppers showed a strong positive correlation between repeat content and genome size. Genome size ranged from 8.2 to 13.7 Gb in these six species with a repeat content ranging routinely from 79% to 87%, with the exception of *Stauroderus scalaris* whose genome is 96% repetitive DNA and the second-largest insect genome documented. Our estimation of genome size for *F. auricularia* doesn’t show gigantism (~968.22Mb, flow cytometer estimate). However, its repeatome (64.62%), is almost twice that of other hemimetabolous insects like *Gryllus bimaculatus* (33.69%) and *Laupala kohalensis* (35.51%) (Ylla *et al.* 2021). This fold increase in the repeatome is surprising given both *G. bimaculatus* and *L. kohalensis* have bigger genomes (~1.6 Gb) than *F. auricularia*.

**Table 2.**
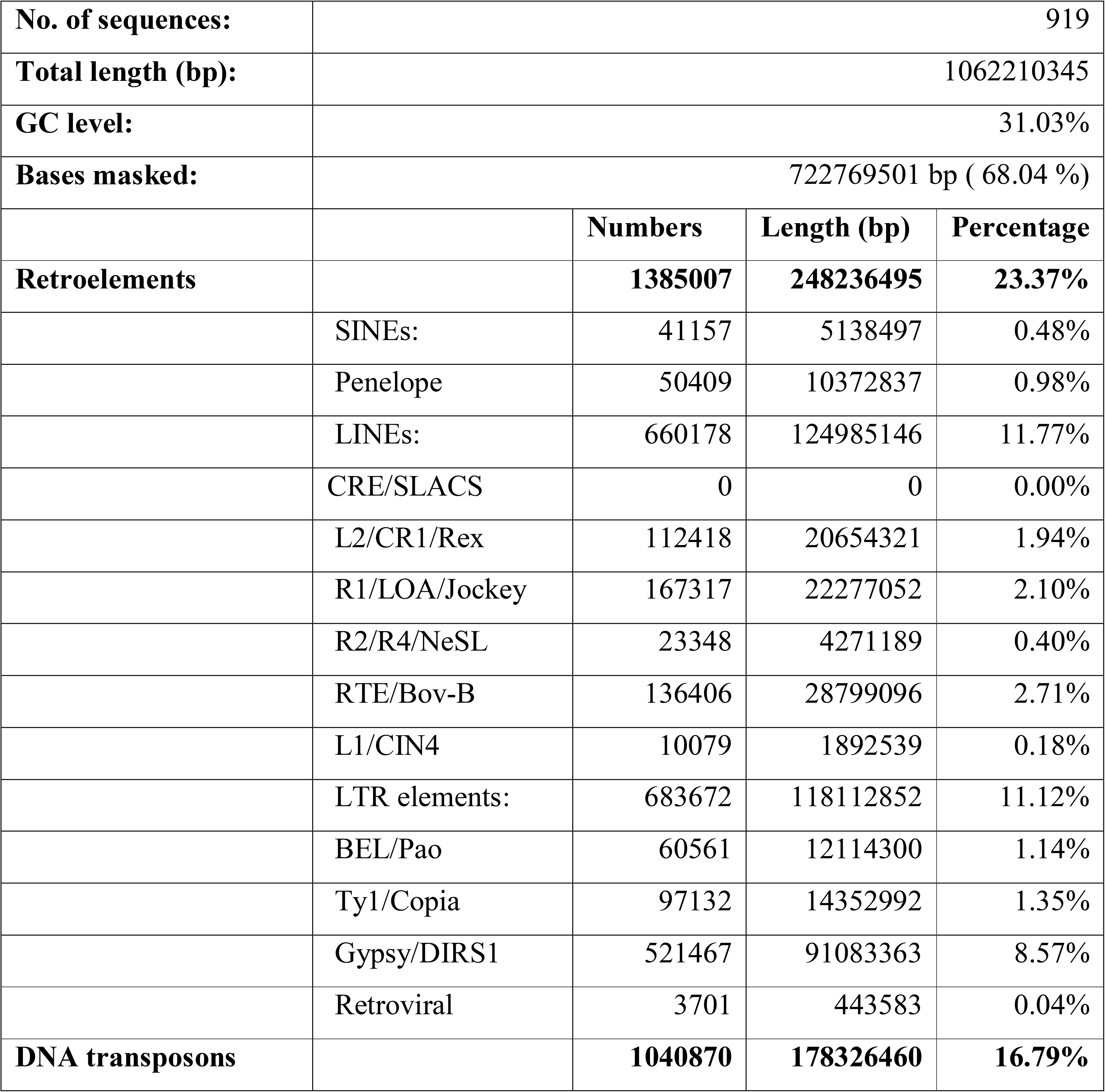
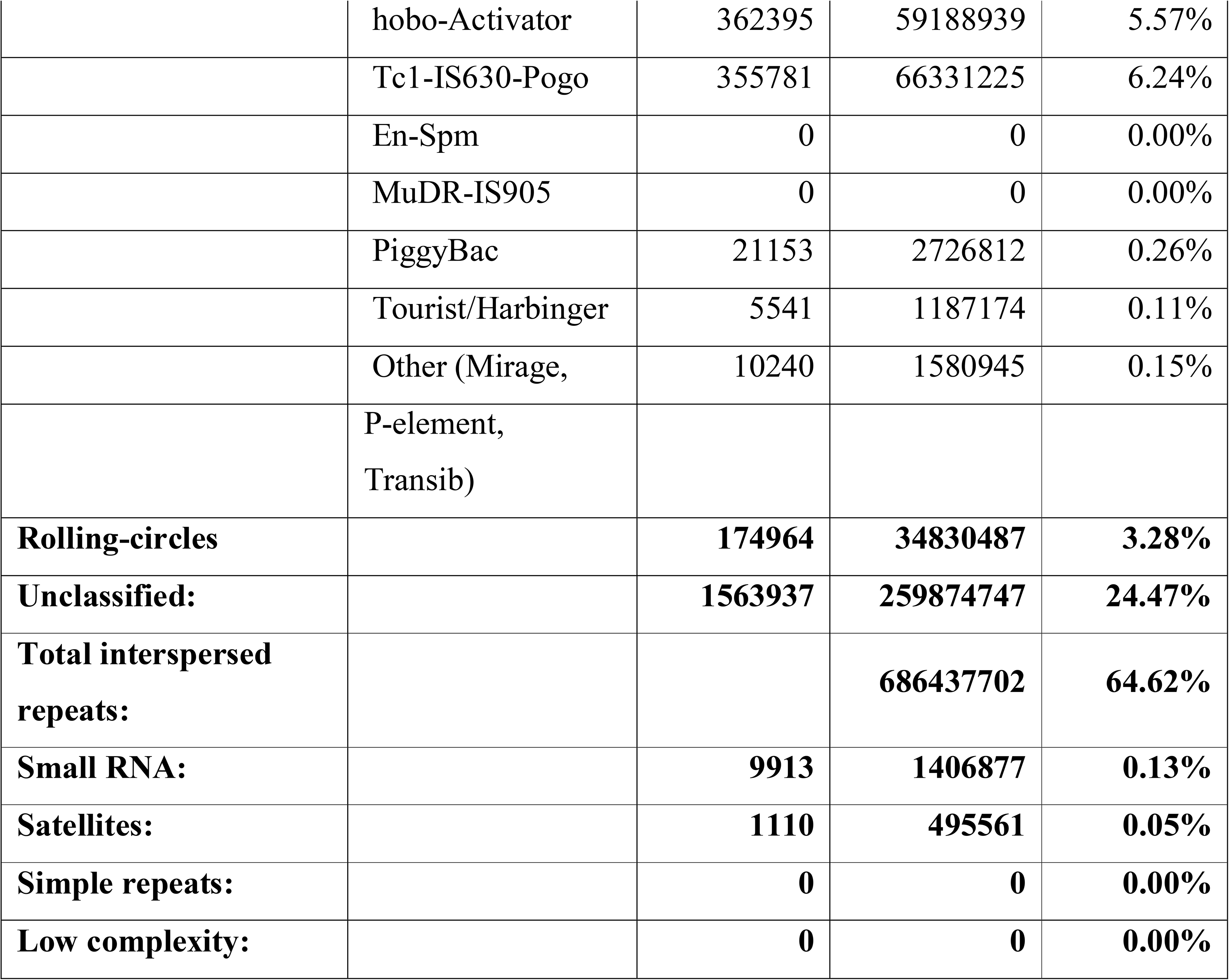
Repeat content analysis in the European earwig *Forficula auricularia* genome.

|  |  |  |  |  |
| --- | --- | --- | --- | --- |
| **No. of sequences:** |  | 919 |  |  |
| **Total length (bp):** |  | 1062210345 |  |  |
| **GC level:** |  | 31.03% |  |  |
| **Bases masked:** |  | 722769501 bp ( 68.04 %) |  |  |
|  |  | **Numbers** | **Length (bp)** | **Percentage** |
| **Retroelements** |  | **1385007** | **248236495** | **23.37%** |
|  | SINEs: | 41157 | 5138497 | 0.48% |
|  | Penelope | 50409 | 10372837 | 0.98% |
|  | LINEs: | 660178 | 124985146 | 11.77% |
|  | CRE/SLACS | 0 | 0 | 0.00% |
|  | L2/CR1/Rex | 112418 | 20654321 | 1.94% |
|  | R1/LOA/Jockey | 167317 | 22277052 | 2.10% |
|  | R2/R4/NeSL | 23348 | 4271189 | 0.40% |
|  | RTE/Bov-B | 136406 | 28799096 | 2.71% |
|  | L1/CIN4 | 10079 | 1892539 | 0.18% |
|  | LTR elements: | 683672 | 118112852 | 11.12% |
|  | BEL/Pao | 60561 | 12114300 | 1.14% |
|  | Ty1/Copia | 97132 | 14352992 | 1.35% |
|  | Gypsy/DIRS1 | 521467 | 91083363 | 8.57% |
|  | Retroviral | 3701 | 443583 | 0.04% |
| **DNA transposons** |  | **1040870** | **178326460** | **16.79%** |
|  | hobo-Activator | 362395 | 59188939 | 5.57% |
|  | Tc1-IS630-Pogo | 355781 | 66331225 | 6.24% |
|  | En-Spm | 0 | 0 | 0.00% |
|  | MuDR-IS905 | 0 | 0 | 0.00% |
|  | PiggyBac | 21153 | 2726812 | 0.26% |
|  | Tourist/Harbinger | 5541 | 1187174 | 0.11% |
|  | Other (Mirage,<br>P-element,<br>Transib) | 10240 | 1580945 | 0.15% |
| **Rolling-circles** |  | **174964** | **34830487** | **3.28%** |
| **Unclassified:** |  | **1563937** | **259874747** | **24.47%** |
| **Total interspersed repeats:** |  |  | **686437702** | **64.62%** |
| **Small RNA:** |  | **9913** | **1406877** | **0.13%** |
| **Satellites:** |  | **1110** | **495561** | **0.05%** |
| **Simple repeats:** |  | **0** | **0** | **0.00%** |
| **Low complexity:** |  | **0** | **0** | **0.00%** |

### Genome annotation

Combining evidence-based and *ab initio* gene models in the MAKER2 pipeline, we identified 12,876 genes and 21,031 mRNAs in the genome assembly. The mean gene length is 12,096 bp and the total gene length is 155.75 Mb, which makes 14.7% of the whole assembly. The longest gene annotated is 412,198 bp and the longest CDS is 19,035 bp (Table 3). 61.35% of total predicted mRNAs and 59.53% of predicted proteins are also functionally annotated through either one or more of InterPro, Gene ontology and Pfam databases (Supplementary Table 3). Comparing with the Arthropoda database we got 74.4% complete BUSCO score for annotated Transcriptome and 70.9% complete BUSCO score for annotated Proteins (Figure 3). 98.3% of the gene models have AED score of 0.5 or less, assuring highly confident gene prediction (Supplementary Figure 1).

**Table 3.**
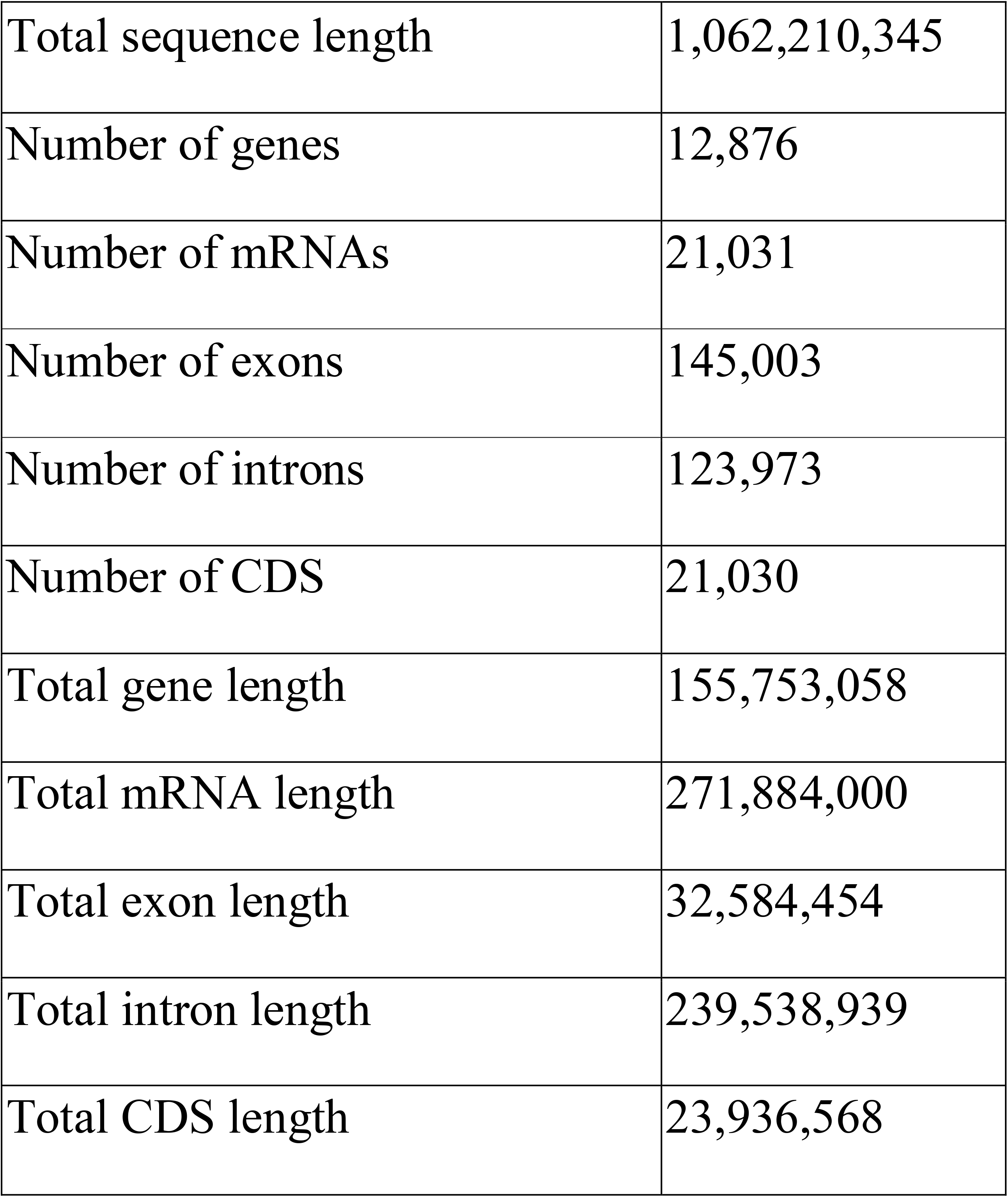
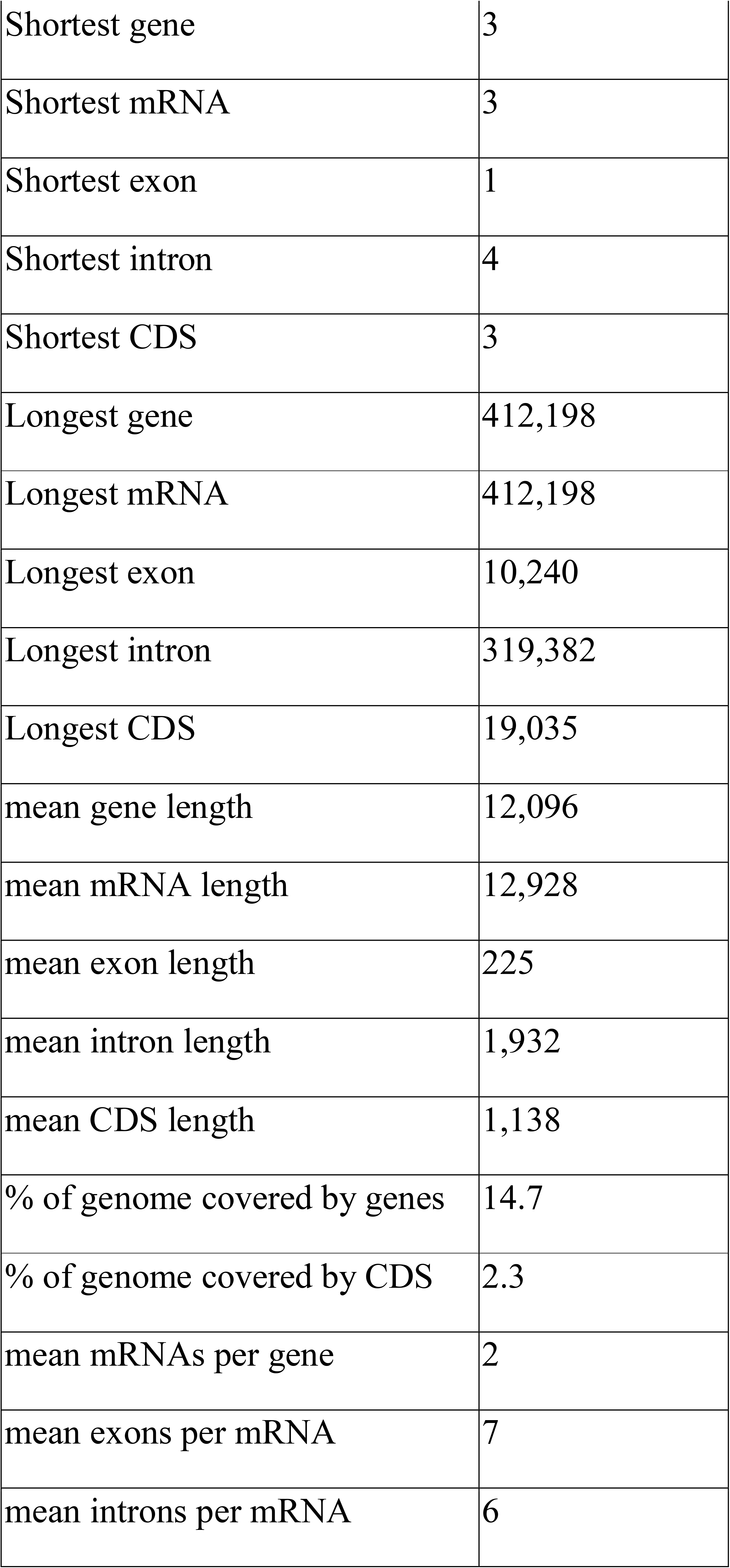
Genome annotation summary for the European earwig *Forficula auricularia*

|  |  |
| --- | --- |
| Total sequence length | 1,062,210,345 |
| Number of genes | 12,876 |
| Number of mRNAs | 21,031 |
| Number of exons | 145,003 |
| Number of introns | 123,973 |
| Number of CDS | 21,030 |
| Total gene length | 155,753,058 |
| Total mRNA length | 271,884,000 |
| Total exon length | 32,584,454 |
| Total intron length | 239,538,939 |
| Total CDS length | 23,936,568 |
| Shortest gene | 3 |
| Shortest mRNA | 3 |
| Shortest exon | 1 |
| Shortest intron | 4 |
| Shortest CDS | 3 |
| Longest gene | 412,198 |
| Longest mRNA | 412,198 |
| Longest exon | 10,240 |
| Longest intron | 319,382 |
| Longest CDS | 19,035 |
| mean gene length | 12,096 |
| mean mRNA length | 12,928 |
| mean exon length | 225 |
| mean intron length | 1,932 |
| mean CDS length | 1,138 |
| % of genome covered by genes | 14.7 |
| % of genome covered by CDS | 2.3 |
| mean mRNAs per gene | 2 |
| mean exons per mRNA | 7 |
| mean introns per mRNA | 6 |

**Figure 3.**
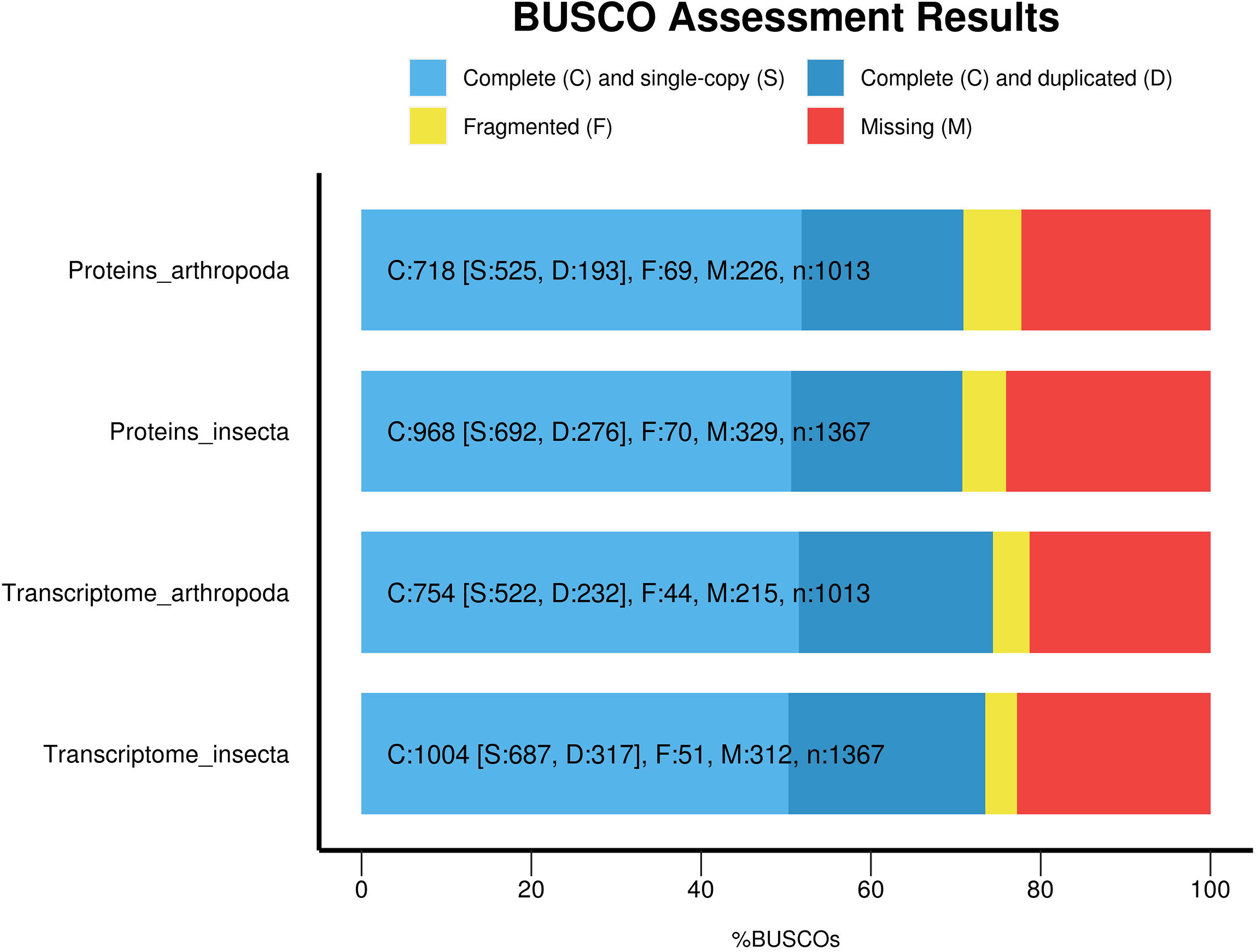
Annotation completeness through BUSCO database. The plot shows the BUSCO (v.5) percentage (x-axis) for the annotated Proteins and Transcriptomes using Arthropoda_odb10 and Insecta_odb10 database as indicated in the y-axis.

The GC content of the *F. auricularia* genome is 31.03%, markedly greater than the 19.3% GC observed in the genome of the earwig *A. maritima* (Mei *et al.* 2022). A comparison of GC content for different genomic regions of *F. auricularia* showed that exons have higher GC content (0.372±0.087) (Mean±SD) and introns have lower (0.267±0.075) when compared between intergenic regions (N = 823,037), genes (N = 12,876), exons (N = 145,003), introns (N = 123,973), and non-overlapping 10kb windows throughout the genome (N = 106,686) (Figure 4). GC content for 10kb windows was 0.308±0.032 which resembles the mean GC content of the whole genome (0.310). This finding is not unexpected as a higher GC content in exons versus introns is common across the animal and plant kingdom because of the evolutionary selection of exon regions (Amit *et al.* 2012). There was a significant different for each pairwise comparison using ANOVA followed by Tukey HSD with *P* < 0.0001.

**Figure 4.**
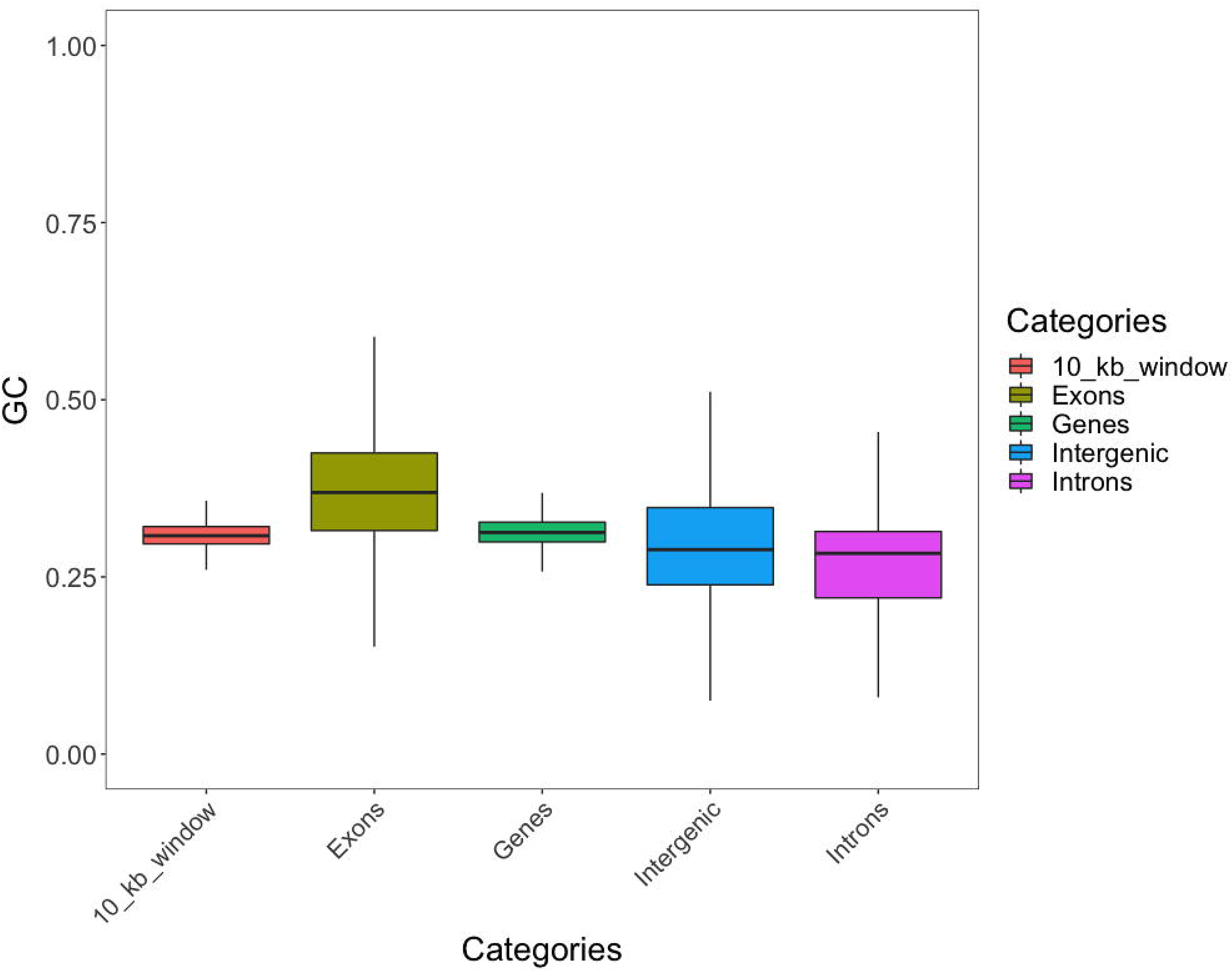
GC percentage in different genomic regions in *F. auricularia* genome. GC content for 10kb windows was generated without regard to any genomic features. Whiskers extend to 25th and 75th percentiles. GC content in exons is higher and in introns is lower compared to the genome average.

In conclusion, the assembled and annotated *F. auricularia* genome presented here is a key resource to develop this important insect species as a genetic model. We anticipate this will enhance the genetic study on various aspects of its biology, including developmental biology, sociality, and evolutionary characteristics.

### Phylogenetic analysis

The phylogenetic analysis showed two distinct subspecies groups within the *F. auricularia* (Figure 5). The two subspecies A and B inferred in our analysis is in complete agreement with the analysis by Wirth et al. (1998). One clade includes 24 individuals including 9 originally identified by Wirth and colleagues as Species A (green circle labels, Figure 5). While the remaining 11 individuals cluster into a separate clade that includes our isolate (red square label, Figure 5) and 6 individuals originally identified by Wirth and colleagues as Species B (green square labels, Figure 5. The analysis confirmed that the isolate sequenced and reported in this paper (Dunedin, NZ) belongs to the Species B of *F. auricularia* species. This is also in accordance with the report from Quarrel et al. (2018). The phylogenetic analysis carried by Quarrel and colleagues reported all Australasian (including two isolates from the New Zealand population) earwig populations as species B lineage.

**Figure 5.**
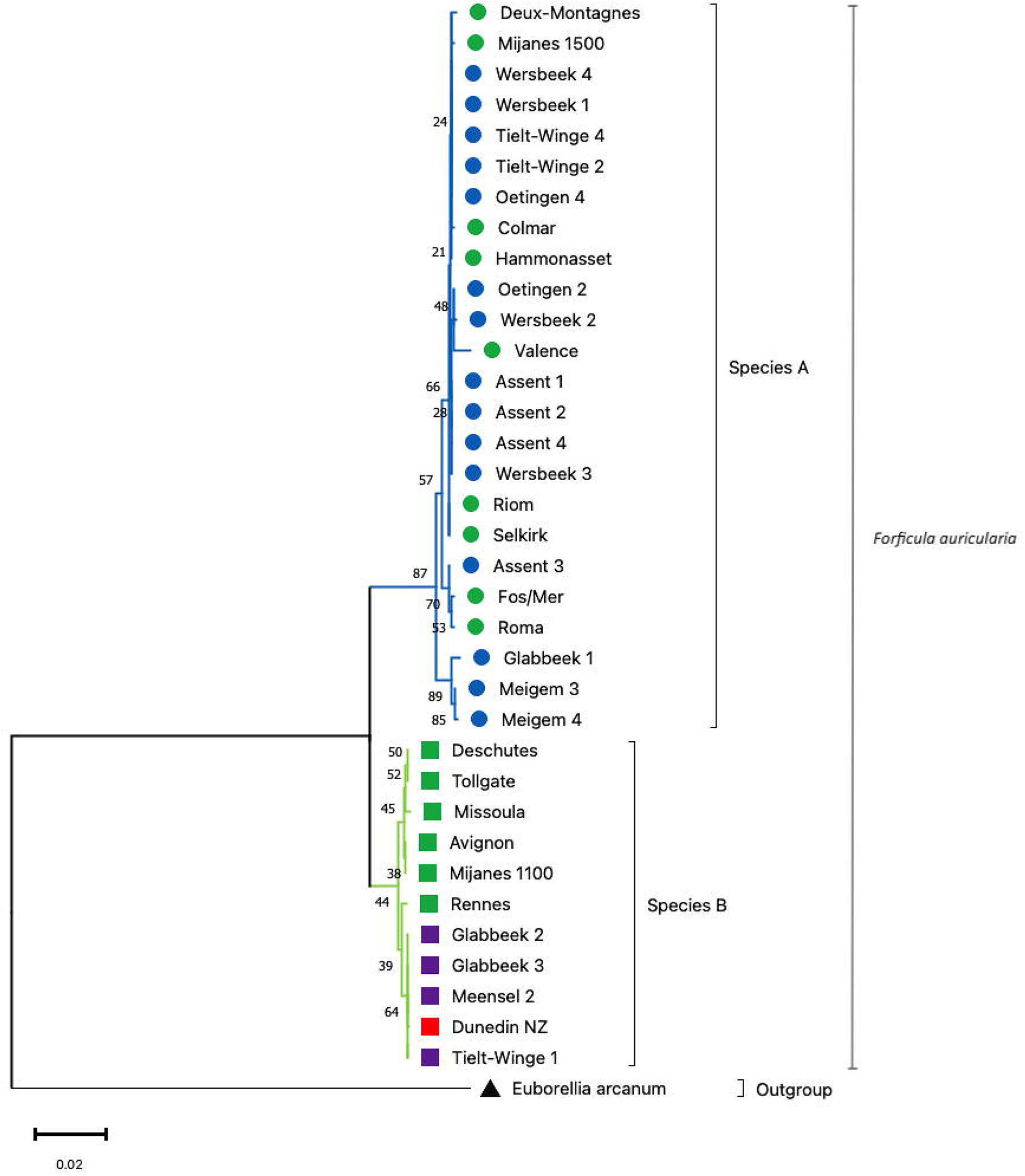
The phylogenetic relationships of different isolates of *Forficula auricularia* inferred by Neighbour-Joining method and Maximum Composite Likelihood approaches using MEGA11. All ambiguous positions were removed for each nucleotide sequence pair (pairwise deletion). The percentage of replicate trees in which the associated taxa clustered together in the bootstrap test (1000 replicates) are shown next to the branches. The tree is drawn to scale, with branch lengths in the same units as those of the evolutionary distances used to infer the phylogenetic tree. Isolates labelled with the coloured squares are Species B. The red square (Dunedin NZ) is the isolate for which the genome is reported in this paper. Green squares are the species categorized as Species B by Wirth et al. (1998) and the purple squares are other isolates for which the nucleotide sequences were downloaded from NCBI. Similarly isolates labelled with coloured circles belong to Species A. Green circles represent isolates inferred as Species A by Wirth et al. (1998) and blue are other isolates for which nucleotide sequences were downloaded from NCBI. *Euborellia arcanum* is the outgroup labelled with a bold triangle.

## Data Availability Statement

The genome assembly and annotation of *Forficula auricularia* are available through FigShare https://doi.org/10.6084/m9.figshare.19092044. The raw sequencing reads are deposited in NCBI with accession number PRJNA800435.

## Code availability

The scripts used for genome assembly, repeat library preparation and masking, and genome annotation, are available at GitHub under GNU GPLv3 license. (https://github.com/upendrabhattarai/Earwig_genome_project)

## Acknowledgements

We thank the support team at New Zealand eScience Infrastructure (NeSI) for their help running various software on their HPC platforms and the staff at Dunedin Botanical Garden, Dunedin, New Zealand, for their help during earwig collection in the field.

## Literature Cited

Alonge, M., L. Lebeigle, M. Kirsche, S. Aganezov, X. Wang et al., 2021 Automated assembly scaffolding elevates a new tomato system for high-throughput genome editing. bioRxiv 2021.11.18.469135.

Amit, M., M. Donyo, D. Hollander, A. Goren, E. Kim et al., 2012 Differential GC Content between Exons and Introns Establishes Distinct Strategies of Splice-Site Recognition. Cell Rep. 1: 543–556.

Binns, M., A. A. Hoffmann, M. Helden, T. Heddle, M. P Hill et al., 2021 Lifecycle of the invasive omnivore, *Forficula auricularia*, in Australian grain growing environments. Pest Manag. Sci. 77: 1818–1828.

Brian Haas, 2010 TransposonPSI: An Application of PSI-Blast to Mine (Retro-)Transposon ORF Homologies.

Campos, M. R., M. C Picanço, J. C. Martins, A. C. Tomaz, and R. N. C. Guedes, 2011 Insecticide selectivity and behavioral response of the earwig *Doru luteipes*. Crop Prot. 30: 1535–1540.

Chakraborty, M., J. G Baldwin-Brown, A. D. Long, and J. J. Emerson, 2016 Contiguous and accurate de novo assembly of metazoan genomes with modest long read coverage. Nucleic Acids Res. 44: gkw654.

Chen, N., 2004 Using RepeatMasker to Identify Repetitive Elements in Genomic Sequences. Curr. Protoc. Bioinforma. 5: 4.10.1–4.10.14.

Coombe, L., J. Zhang, B. P. Vandervalk, J. Chu, S. D Jackman et al., 2018 ARKS: chromosome-scale scaffolding of human genome drafts with linked read kmers. BMC Bioinformatics 19: 234.

Edgar, R. C., 2010 Search and clustering orders of magnitude faster than BLAST. Bioinformatics 26: 2460–2461.

Ellinghaus, D., S. Kurtz, and U. Willhoeft, 2008 LTRharvest, an efficient and flexible software for de novo detection of LTR retrotransposons. BMC Bioinformatics 9: 18.

Falk, J., J. W. Y. Wong, M. Kölliker, and J. Meunier, 2014 Sibling Cooperation in Earwig Families Provides Insights into the Early Evolution of Social Life. Am. Nat. 183: 547–557.

Felsenstein, J., 1985 Confidence Limits on Phylogenies: An Approach Using the Bootstrap. Evolution (N. Y). 39: 783.

Flynn, J. M., R. Hubley, C. Goubert, J. Rosen, A. G Clark et al., 2020 RepeatModeler2 for automated genomic discovery of transposable element families. Proc. Natl. Acad. Sci. 117: 9451–9457.

Forslund, P., 2000 Male–male competition and large size mating advantage in European earwigs, *Forficula auricularia*. Anim. Behav. 59: 753–762.

Grabherr, M. G., B. J. Haas, M. Yassour, J. Z. Levin, D. A Thompson et al., 2011 Full-length transcriptome assembly from RNA-Seq data without a reference genome. Nat. Biotechnol. 29: 644–652.

Gregory, T. R., 2005 Genome Size Evolution in Animals, pp. 3–87 in The Evolution of the Genome, edited by T. R. B. T.-T. E. of the G. Gregory. Elsevier, Burlington.

Gurevich, A., V. Saveliev, N. Vyahhi, and G. Tesler, 2013 QUAST: quality assessment tool for genome assemblies. Bioinformatics 29: 1072–1075.

Herbison, R. E. H., S. Evans, J.-F. Doherty, and R. Poulin, 2019 Let’s go swimming: mermithid-infected earwigs exhibit positive hydrotaxis. Parasitology 146: 1631–1635.

Hill, M. P., M. Binns, P. A. Umina, A. A. Hoffmann, and S. Macfadyen, 2019 Climate, human influence and the distribution limits of the invasive European earwig, *Forficula auricularia*, in Australia. Pest Manag. Sci. 75: 134–143.

Hiltunen, M., M. Ryberg, and H. Johannesson, 2021 ARBitR: an overlap-aware genome assembly scaffolder for linked reads (M. Pier Luigi, Ed.). Bioinformatics 37: 2203–2205.

Hoff, K. J., A. Lomsadze, M. Borodovsky, and M. Stanke, 2019 Whole-Genome Annotation with BRAKER, pp. 65–95 in Methods in molecular biology (Clifton, N.J.),.

Holt, C., and M. Yandell, 2011 MAKER2: an annotation pipeline and genome-database management tool for second-generation genome projects. BMC Bioinformatics 12: 491.

Jones, P., D. Binns, H.-Y. Chang, M. Fraser, W. Li et al., 2014 InterProScan 5: genome-scale protein function classification. Bioinformatics 30: 1236–1240.

Kamimura, Y., 2006 Right-handed penises of the earwig *Labidura riparia* (Insecta, Dermaptera, Labiduridae): Evolutionary relationships between structural and behavioral asymmetries. J. Morphol. 267: 1381–1389.

Kamimura, Y., Y. Matsumura, C.-C. S. Yang, and S. N. Gorb, 2021 Random or handedness? Use of laterally paired penises in Nala earwigs (Insecta: Dermaptera: Labiduridae). Biol. J. Linn. Soc. 134: 716–731.

Kokot, M., M. Długosz, and S. Deorowicz, 2017 KMC 3: counting and manipulating k-mer statistics (B. Berger, Ed.). Bioinformatics 33: 2759–2761.

Kolmogorov, M., J. Yuan, Y. Lin, and P. A. Pevzner, 2019 Assembly of long, error-prone reads using repeat graphs. Nat. Biotechnol. 37: 540–546.

Kramer, J., J. Thesing, and J. Meunier, 2015 Negative association between parental care and sibling cooperation in earwigs: a new perspective on the early evolution of family life? J. Evol. Biol. 28: 1299–1308.

Labandeira, C. C., and T. L. Phillips, 1996 A Carboniferous insect gall: insight into early ecologic history of the Holometabola. Proc. Natl. Acad. Sci. 93: 8470–8474.

Laetsch, D. R., and M. L. Blaxter, 2017 BlobTools: Interrogation of genome assemblies. F1000Research 6: 1287.

Mei, Y., D. Jing, S. Tang, X. Chen, H. Chen et al., 2022 InsectBase 2.0: a comprehensive gene resource for insects. Nucleic Acids Res. 50: D1040–D1045.

Van Meyel, S., and J. Meunier, 2020 Filial egg cannibalism in the European earwig: its determinants and implications in the evolution of maternal egg care. Anim. Behav. 164: 155–162.

Mukherjee, K., R. M. Twyman, and A. Vilcinskas, 2015 Insects as models to study the epigenetic basis of disease. Prog. Biophys. Mol. Biol. 118: 69–78.

Qin, M., S. Wu, A. Li, F. Zhao, H. Feng et al., 2019 LRScaf: improving draft genomes using long noisy reads. BMC Genomics 20: 955.

Quarrell, S. R., J. Arabi, A. Suwalski, M. Veuille, T. Wirth et al., 2018 The invasion biology of the invasive earwig, *Forficula auricularia* in Australasian ecosystems. Biol. Invasions 20: 1553–1565.

Roach, M. J., S. A. Schmidt, and A. R. Borneman, 2018 Purge Haplotigs: allelic contig reassignment for third-gen diploid genome assemblies. BMC Bioinformatics 19: 460.

Saitou, N., and M. Nei, 1987 The neighbor-joining method: a new method for reconstructing phylogenetic trees. Mol. Biol. Evol. 4: 406–425.

Simão, F. A., R. M. Waterhouse, P. Ioannidis, E. V. Kriventseva, and E. M. Zdobnov, 2015 BUSCO: assessing genome assembly and annotation completeness with single-copy orthologs. Bioinformatics 31: 3210–3212.

Simons, M., and E. Tibbetts, 2019 Insects as models for studying the evolution of animal cognition. Curr. Opin. Insect Sci. 34: 117–122.

Song, L., D. S. Shankar, and L. Florea, 2016 Rascaf: Improving Genome Assembly with RNA Sequencing Data. Plant Genome 9:.

Steinbiss, S., U. Willhoeft, G. Gremme, and S. Kurtz, 2009 Fine-grained annotation and classification of de novo predicted LTR retrotransposons. Nucleic Acids Res. 37: 7002–7013.

Tamura, K., M. Nei, and S. Kumar, 2004 Prospects for inferring very large phylogenies by using the neighbor-joining method. Proc. Natl. Acad. Sci. 101: 11030–11035.

Tamura, K., G. Stecher, and S. Kumar, 2021 MEGA11: Molecular Evolutionary Genetics Analysis Version 11 (F. U. Battistuzzi, Ed.). Mol. Biol. Evol. 38: 3022–3027.

Tourneur, J.-C., and J. Meunier, 2019 Thermal regimes, but not mean temperatures, drive patterns of rapid climate adaptation at a continent-scale: evidence from the introduced European earwig across North America. bioRxiv 550319.

Truman, J. W., 2019 The Evolution of Insect Metamorphosis. Curr. Biol. 29: R1252–R1268.

Vassetzky, N. S., and D. A. Kramerov, 2013 SINEBase: a database and tool for SINE analysis. Nucleic Acids Res. 41: D83–D89.

Warren, R. L., 2016 RAILS and Cobbler: Scaffolding and automated finishing of draft genomes using long DNA sequences. J. Open Source Softw. 1: 116.

Warren, R. L., C. Yang, B. P. Vandervalk, B. Behsaz, A. Lagman et al., 2015 LINKS: Scalable, alignment-free scaffolding of draft genomes with long reads. Gigascience 4: 35.

Weisenfeld, N. I., V. Kumar, P. Shah, D. M. Church, and D. B. Jaffe, 2017 Direct determination of diploid genome sequences. Genome Res. 27: 757–767.

Wilson-Sanders, S. E., 2011 Invertebrate Models for Biomedical Research, Testing, and Education. ILAR J. 52: 126–152.

Winnebeck, E. C., C. D. Millar, and G. R. Warman, 2010 Why Does Insect RNA Look Degraded? J. Insect Sci. 10: 1–7.

Wirth, T., R. Le Guellec, M. Vancassel, and M. Veuille, 1998 Molecular and Reproductive Characterization of Sibling Species in the European Earwig (Forficula auricularia). Evolution (N. Y). 52: 260.

Wu, C., V. G. Twort, R. N. Crowhurst, R. D. Newcomb, and T. R. Buckley, 2017 Assembling large genomes: analysis of the stick insect (*Clitarchus hookeri*) genome reveals a high repeat content and sex-biased genes associated with reproduction. BMC Genomics 18: 884.

Xu, G.-C., T.-J. Xu, R. Zhu, Y. Zhang, S.-Q. Li et al., 2019 LR_Gapcloser: a tiling path-based gap closer that uses long reads to complete genome assembly. Gigascience 8: giy157

Yang, A. S., 2001 Modularity, evolvability, and adaptive radiations: a comparison of the hemi-and holometabolous insects. Evol. Dev. 3: 59–72.

Ylla, G., T. Nakamura, T. Itoh, R. Kajitani, A. Toyoda et al., 2021 Insights into the genomic evolution of insects from cricket genomes. Commun. Biol. 4: 733.

