## Supplementary Table 1, Supplementary Table 2, Supplementary Table 3, Supplementary Figure 1 for "A high-quality genome assembly and annotation of the European earwig *Forficula auricularia*" (from another version of this preprint: version 1, 2022-02-02)

[*] Corresponding authors

**Supplementary Table 1**. Nanopore sequencing output

| Number of Reads | Total Bases | Median Read Length | N50 Length | Median Read Quality |
|---|---|---|---|---|
| 3,021,511.00 | 10,698,430,965.00 | 897.00 | 11,986.00 | 13.34 |

**Supplementary Table 2**. Assembly statistics after each round of processing

| Assembly steps | Assembly length | No. of scaffolds | N50 | L50 | Ns per 100 kbp | Busco % (Quast) | |
|---|---|---|---|---|---|---|---|
| | | | | | | Complete | Partial |
| Supernova | 1,145,470,221 | 145,055 | 30,358 | 7,500 | 3,677.89 | 64.69 | 9.24 |
| Flye | 1,118,374,848 | 18,766 | 180,737 | 1,832 | 0.35 | 82.18 | 9.24 |
| Quickmerge | 1138492550 | 16670 | 222450 | 1435 | 1028.63 | 84.16 | 5.94 |
| Purgehaplotigs | 1092680447 | 11914 | 233917 | 1342 | 1053.46 | 84.49 | 5.94 |
| Rails & Cobbler | 1094435828 | 10883 | 251120 | 1250 | 851.82 | 84.82 | 5.61 |
| Lrscaff | 1213856231 | 8227 | 329347 | 1101 | 2060.89 | 84.49 | 4.95 |
| LrGapcloser | 1213833977 | 8227 | 329347 | 1101 | 488.12 | 86.8 | 3.96 |
| Ragtag | 1213843077 | 8136 | 330338 | 1097 | 488.84 | 87.79 | 3.63 |
| Arbit | 1211594836 | 6512 | 529318 | 681 | 500.51 | 87.13 | 3.63 |
| Arks & Links | 1211713036 | 5330 | 740184 | 473 | 510.23 | - | - |
| Rascaf | 1220673574 | 5170 | 824726 | 410 | 1241.57 | 87.13 | 3.96 |

| | | | | | | | |
|---|---|---|---|---|---|---|---|
| Purgehaplotigs | 1134953563 | 4307 | 887017 | 363 | 864.02 | 85.81 | 4.62 |
| Ragtag | 1135017063 | 3672 | 1234415 | 225 | 869.52 | 86.8 | 4.29 |
| Blobtools | 1062122003 | 1881 | 1349000 | 199 | 837.87 | 83.17 | 4.62 |
| Ragtag | 1062218203 | 919 | 12548662 | 20 | 846.85 | 83.17 | 4.62 |
| Pilon | 1062210345 | 919 | 12548649 | 20 | 846.85 | 87.13 | 2.97 |

(Note: Missing values are because of no data produced by Quast)

**Supplementary Table 3**. Functional annotation statistics from different databases

| Database | No. of terms linked to mRNA | No. of mRNA with term | No. of gene with term |
|---|---|---|---|
| InterPro | 17976 | 12904 | 7666 |
| Gene ontology term | 15088 | 7669 | 4560 |
| Pfam | 18130 | 12904 | 7666 |

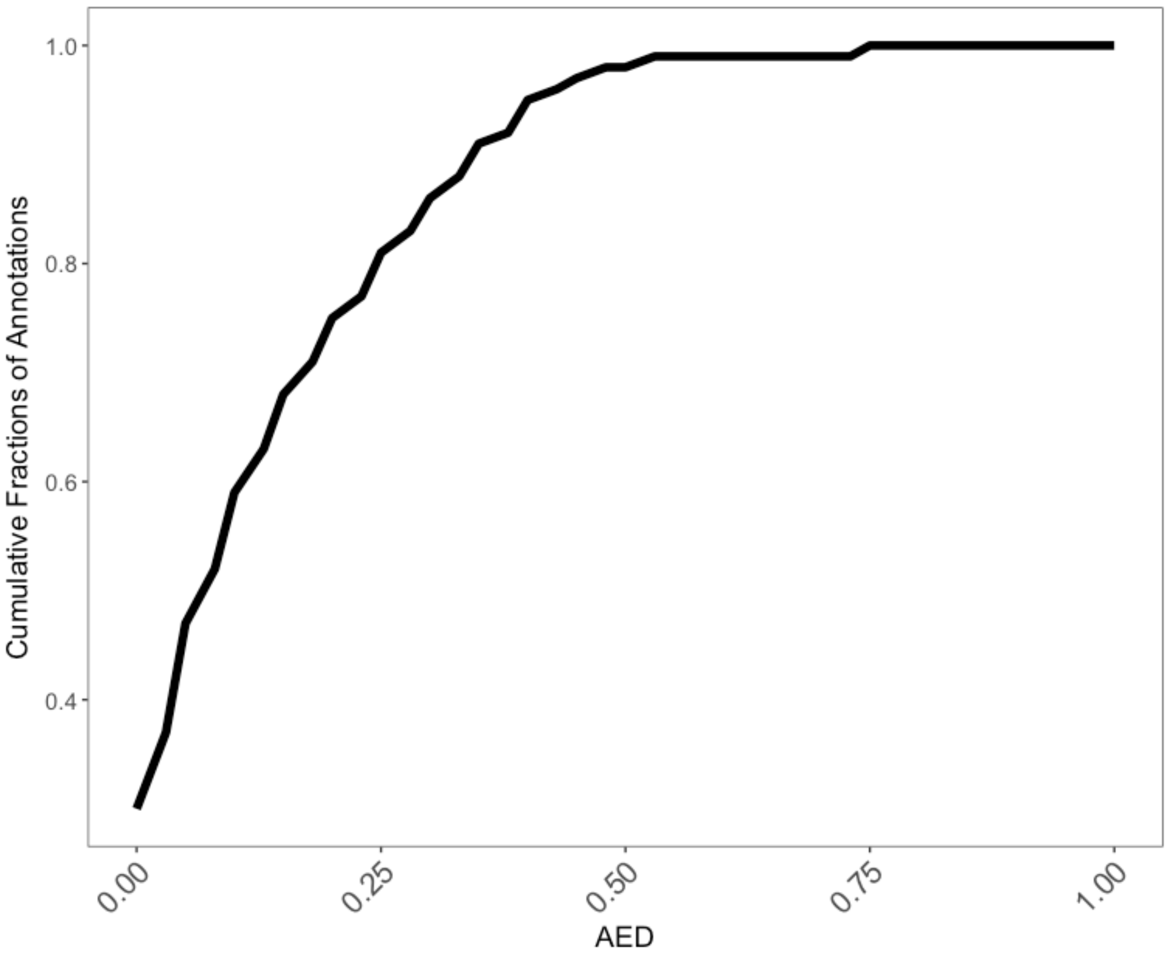


**Supplementary Figure 1**. Annotation quality with AED scores. Y-axis shows the cumulative fractions of annotations and x-axis their AED scores.
